# DNA damage response defects induced by the formation of TDP-43 and mutant FUS cytoplasmic inclusions and their pharmacological rescue

**DOI:** 10.1101/2024.08.22.609115

**Authors:** Stefania Farina, Francesca Esposito, Stefania Modafferi, Ornella Brandi, Michela Di Salvio, Ilaria Della Valle, Giada Cicio, Adelaide Riccardi, Federica Pisati, Anna Garbelli, Nadia D’Ambrosi, Mauro Cozzolino, Gianluca Cestra, Fabrizio d’Adda di Fagagna, Ubaldo Gioia, Sofia Francia

## Abstract

Formation of cytoplasmic inclusions (CIs) of TDP-43 and FUS, along with DNA damage accumulation, is a hallmark of affected motor neurons in Amyotrophic Lateral Sclerosis (ALS). However, the impact of CIs on DNA damage response (DDR) and repair in this pathology remains unprobed. Here, we show that CIs of TDP-43 and FUS^P525L^, co-localizing with stress granules, lead to a dysfunctional DDR activation associated with physical DNA breakage. Inhibition of the activity of the DDR kinase ATM, but not of ATR, abolishes DDR signaling, indicating that DNA double-strand breaks (DSBs) are the primary source of DDR activation. In addition, cells with TDP-43 and FUS^P525L^ CIs exhibit reduced DNA damage-induced RNA synthesis at DSBs. We previously showed that the two endoribonucleases DROSHA and DICER, also known to interact with TDP-43 and FUS during small RNA processing, contribute to DDR signaling at DSBs. Treatment with enoxacin, which stimulates DDR and repair by boosting the enzymatic activity of DICER, restores a proficient DDR and reduces DNA damage accumulation in cultured cells with CIs and *in vivo* in a murine model of ALS. In *Drosophila melanogaster*, Dicer-2 overexpression rescues TDP-43-mediated retinal degeneration. In summary, our results indicate that the harmful effects caused by TDP-43 and FUS proteinopathies include genotoxic stress and that the pharmacological stimulation of the DNA damage signaling and repair counteracts it.

## INTRODUCTION

Amyotrophic Lateral Sclerosis (ALS) is a neurodegenerative disease characterized by the progressive loss of the upper and lower motor neurons due to neuronal cell death^1^. Analogously to other neurodegenerative disorders like Frontotemporal Dementia (FTD) and Alzheimer’s disease (AD), ALS is defined as a proteinopathy since motor neurons of patients affected by this pathology harbor aberrant protein aggregates in their cytoplasm^2^. 90% of ALS cases are sporadic (sALS), thus lacking a familial history, while the remaining ALS cases are characterized by autosomal dominant, autosomal recessive, or X-linked mutations occurring in specific genes, leading to familial inheritance (familial ALS, fALS)^3^. Notably, around 3% of fALS-associated mutations fall in the gene encoding for TAR DNA-binding protein 43 (TDP-43) and 6% in the gene for fused in sarcoma (FUS)^3^. Some mutations, particularly FUS^P525L^, are believed to confer to these proteins an increased predisposition to form cytosolic aggregates^4^. Nevertheless, the vast majority of both sALS and fALS and in all FTD cases, TDP-43 accumulates in CIs in its wild-type form^5^.

TDP-43 and FUS are two RNA-binding proteins (RBPs) that cover multiple functions in the RNA metabolism. For instance, they are both involved in miRNA biogenesis, through their interaction with the endoribonuclease DROSHA^6,7^. Moreover, FUS also exhibits a transcriptional regulatory role and can directly bind to the C-terminal domain (CTD) of RNA polymerase II (RNAPII)^8,9^. Both TDP-43 and FUS are structurally characterized by the presence of low complexity domains (LCDs), which allow the two proteins to undergo liquid-liquid phase separation (LLPS), essential for the formation of insoluble membrane-less organelles, including stress granules (SGs)^10^.

SGs are generated in response to different stress sources, with the proposed function of pausing protein synthesis until the stress ends^11^. Along with their constitutive components, like TIA-1 and G3BP1, SGs may include TDP-43 and FUS^11,12^. Interestingly, TDP-43 depletion alters both G3BP1 and TIA-1 levels and affects SG structure and dynamics^13^. Besides, motor neurons derived from induced pluripotent stem cells (iPSCs) carrying the FUS^P525L^ mutation display an increased number and altered kinetics of SG assembly^14^.

Genome integrity ensures the maintenance of the genetic information against possible unfaithful transmission across generations. Both physiological events and environmental factors can cause DNA breaks, leading to genome instability^15^. Therefore, cells have developed a complex pathway named DNA Damage Response (DDR), which immediately detects the DNA lesion, signals its formation and ensures a proper DNA repair^16^. In particular, following double-strand break (DSB) formation, the multiprotein complex MRN (MRE11-RAD50-NBS1) binds the DNA ends, thus recruiting and activating ATM, a member of the phosphatidylinositol 3-kinase-related kinase (PIKK) family^17^. ATM, in turn, initiates the DDR signaling by phosphorylating the histone variant H2AX, which in its phosphorylated version is referred to as γH2AX^18^. Upon accumulation of single-strand breaks (SSBs), generally occurring during replication stress, H2AX is phosphorylated by ATR, another PIKK^19^. Initial γH2AX chromatin decoration creates a positive feedback loop that fuels further spreading of γH2AX and sequential accumulation of DDR proteins^18^, including 53BP1, which plays a pivotal role in orchestrating the choice of DSB repair pathways^20^.

Different neurodegenerative diseases, including ALS, have been associated with a high accumulation of DNA lesions in the nucleus of motorneurons^21,22^, affecting their viability. Intriguingly, a role for TDP-43 and FUS in DNA repair has been recently identified in TDP-43 and FUS loss of function model systems^23,24^. However, the impact of the formation of TDP-43- and FUS-containing SGs on genome integrity and DDR activation remains unknown.

In the present study, we investigate the molecular mechanisms underlying defects in DDR activation and the consequent accumulation of DNA damage following the formation of TDP-43 and FUS CIs. We also provide evidence that demonstrate that DDR components are potential therapeutic targets to mitigate the neurotoxic effects of TDP-43 and FUS proteinopathies.

## RESULTS

### Formation of TDP-43 and mutant FUS cytoplasmic inclusions (CIs) is associated with DNA damage accumulation

To acutely induce the formation of CIs in human cultured cells, we transiently transfected HeLa cells with plasmids expressing wild-type (WT) FUS and its ALS-linked mutant form (FUS^P525L^), along with WT TDP-43 and two ALS-related mutant derivatives (TDP-43^A382T^ and TDP-43^I383V^), in parallel with an empty vector (EV) as a control (Figure 1 and S1). Western blot analyses confirmed that the expression levels of the mutant isoforms of both TDP-43 and FUS were comparable to their WT counterparts and similar to their endogenous levels (Figure S1A,B). We next probed for FUS and TDP-43 by immunofluorescence (IF) and observed that the overexpression of FUS^P525L^ and WT TDP-43, or its ALS-linked mutants, was associated with CI formation in approximately 30% and 15-20% of the cells, respectively (Figure 1A,B and S1C,D). Differently, the fraction of cells showing endogenous TDP-43-positive CIs was much lower (less than 3%) in samples transfected with the EV or with the plasmid expressing WT FUS (Figure 1A,B and S1C,D). Importantly, FUS segregated to CIs only upon the expression of its mutant version, while it was retained in the nucleus in WT FUS-expressing cells (Figure 1A,B and S1E). Differently, ALS-linked mutations of TDP-43 did not further increase the tendency of this protein to form CIs (Figure 1A,B and S1C,E). This observation aligns with the prevalence of WT TDP-43 inclusions in most ALS patients^5^. Consistently with previous reports^25^, the expression of FUS^P525L^ and TDP-43 induced the formation of SGs (Figure S1E), and FUS^P525L^ and TDP-43 colocalized with such organelles (Figure S1E), as also already observed in ALS brain tissues^25^. By confocal imaging analyses, we noticed that in this setting TDP-43 accumulation in CIs was not accompanied by its nuclear depletion, as others reported^26,27^ (Figure 1A,B and S1E). Conversely, FUS^P525L^ signal was exclusively localized to CIs, with no detectable presence in the nucleus (Figure 1A,B and S1E). These findings suggest a gain-of-function condition for TDP-43 and a combination of gain and loss of function for FUS in our experimental settings. Since the formation of CIs upon expression of either WT or mutant TDP-43 did not exhibit significant differences (Figure 1A,B and S1C,E), and given our research focus on investigating the impact of CI formation on DDR activation and DNA repair, we exclusively conducted experiments using cells expressing WT TDP-43 for the remainder of this study.

**Figure 1.**
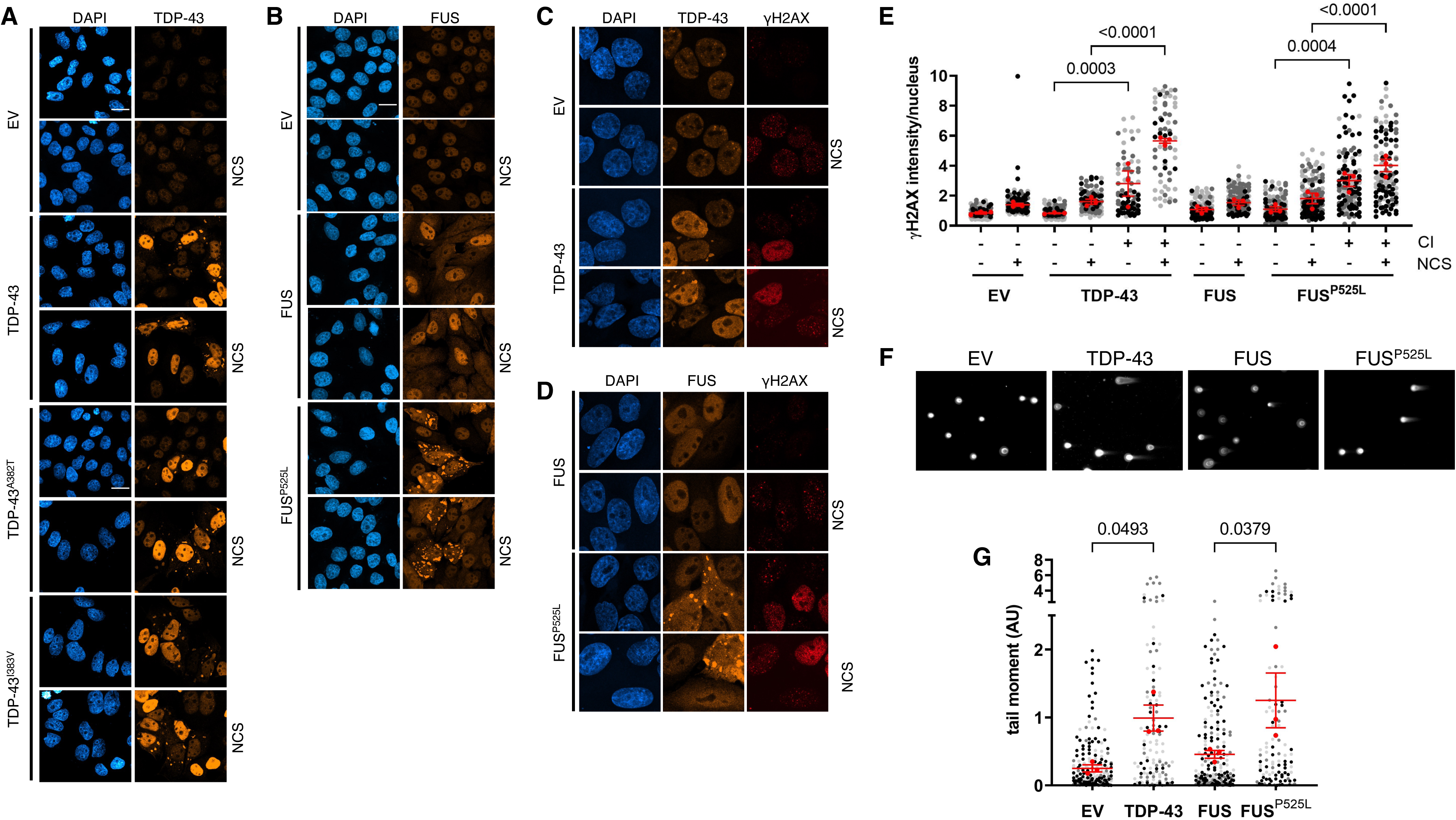
Cells with TDP-43 or FUS^P525L^ cytoplasmic inclusions (CIs) show elevated γH2AX levels and increased DSB formation. **A, B)** Immunofluorescence (IF) analysis of CI formation in damaged (NCS) or undamaged HeLa cells transfected with plasmids expressing wild-type or mutant TDP-43 or FUS; cells transfected with an empty vector (EV) were used as a control; nuclei were counter-stained with DAPI; scale bar = 10 µm. **C, D)** Analysis by IF of γH2AX signals in HeLa cells treated as in A and B. **E)** Super-plot showing the quantification of γH2AX intensity in HeLa cells with or without cytoplasmic inclusions (± CI) determined in C and D. Red dots represent the mean values of each biological replicate; red bars indicate the average ± SEM of three independent experiments. **F)** Representative images of comet assay experiments performed in undamaged HeLa cells transfected as in A and B. **G)** Tail moment analysis of HeLa cells shown in F. The red dots of the super-plot represent the mean values of each biological replicate; red bars indicate the average ± SEM of three independent experiments.

We next investigated the effects of CI formation on DNA damage accumulation at single-cell resolution by treating FUS- or TDP-43-expressing cells with Neocarzinostatin (NCS) – a radiomimetic drug largely used to acutely generate DNA damage^28,29^ – followed by IF analysis. Notably, localization of FUS^P525L^ or TDP-43 into CIs was not affected by NCS administration (Figure S1C,D), indicating that DNA damage is not the cause of FUS^P525L^ or TDP-43 aggregation, but rather a consequence. WT FUS- or EV-transfected cells presented little endogenous DNA damage and properly formed γH2AX-positive DDR foci upon treatment with NCS (Figure 1C-E). Differently, a robust accumulation of pan-nuclear γH2AX signal was observed in cells with mutant FUS and TDP-43 CIs (Figure 1C-E). Such a massive amount of γH2AX occurred even in the absence of exogenously inflicted DNA damage (Figure 1C-E), indicating that the formation of CIs following FUS^P525L^ or TDP-43 expression was sufficient to induce a significant genotoxic stress. This intense γH2AX signal prevents from distinguishing discrete γH2AX foci in NCS-treated cells (Figure 1C-E). Importantly, cells without mutant FUS- or TDP-43 CIs did not display pan-nuclear γH2AX signal, both in undamaged and damaged conditions (Figure 1C-E). These results indicate that genotoxicity is rapidly and specifically induced upon the formation of FUS^P525L^ or TDP-43 positive SGs.

γH2AX accumulation is indicative of an increase of unrepaired DSBs. Thus, to specifically detect DSB accumulation following FUS or TDP-43 expression, we performed a neutral comet assay in cells not exposed to exogenous DNA damage and transfected with EV, FUS, FUS^P525L^ or TDP-43. Interestingly, we observed that the expression of TDP-43 or FUS^P525L^ increased the comet tail moment, suggestive of a higher amount of DSBs, compared to EV-transfected samples or cells overexpressing WT FUS (Figure 1F,G).

These results indicate that the elevated γH2AX signal observed in the nucleus of cells with mutant FUS- or TDP-43-positive CIs is associated with a significant increase of physical DNA breakage in the form of DSBs.

### ATM is hyper-activated and responsible for the increased **γ**H2AX levels in cells with CIs

As soon as the DNA damage is sensed, the apical kinases ATM and ATR are known to be engaged and trigger the local phosphorylation of H2AX, along with several other protein targets^18,19^. A previous study showed that pan-nuclear γH2AX accumulates in response to ATM activation in different contexts of chromatin structure alteration^30^. Instead, ATR activation has been occasionally associated with γH2AX pan-nuclear signal during replication stress^31,32^. Thus, to determine which of these two kinases was responsible for γH2AX accumulation observed in cells bearing CIs, we treated cells expressing FUS^P525L^, TDP-43, or control samples, with specific ATM or ATR kinase activity inhibitors (KU60019 or VE-821, respectively^33^) and monitored their impact on γH2AX signals by IF. The specificity of inhibition of the two DDR kinases was confirmed by monitoring their auto-phosphorylation through immunoblotting (Figure S2A). Intriguingly, cells with TDP-43 or FUS^P525L^ CIs showed a significant reduction of γH2AX signal upon ATM inhibition, compared to control samples treated with the vehicle (DMSO) (Figure 2A,B), while, ATR inactivation had a little impact in reducing γH2AX levels in cells with FUS^P525L^ or TDP-43 CIs (Figure 2A,B).

**Figure 2.**
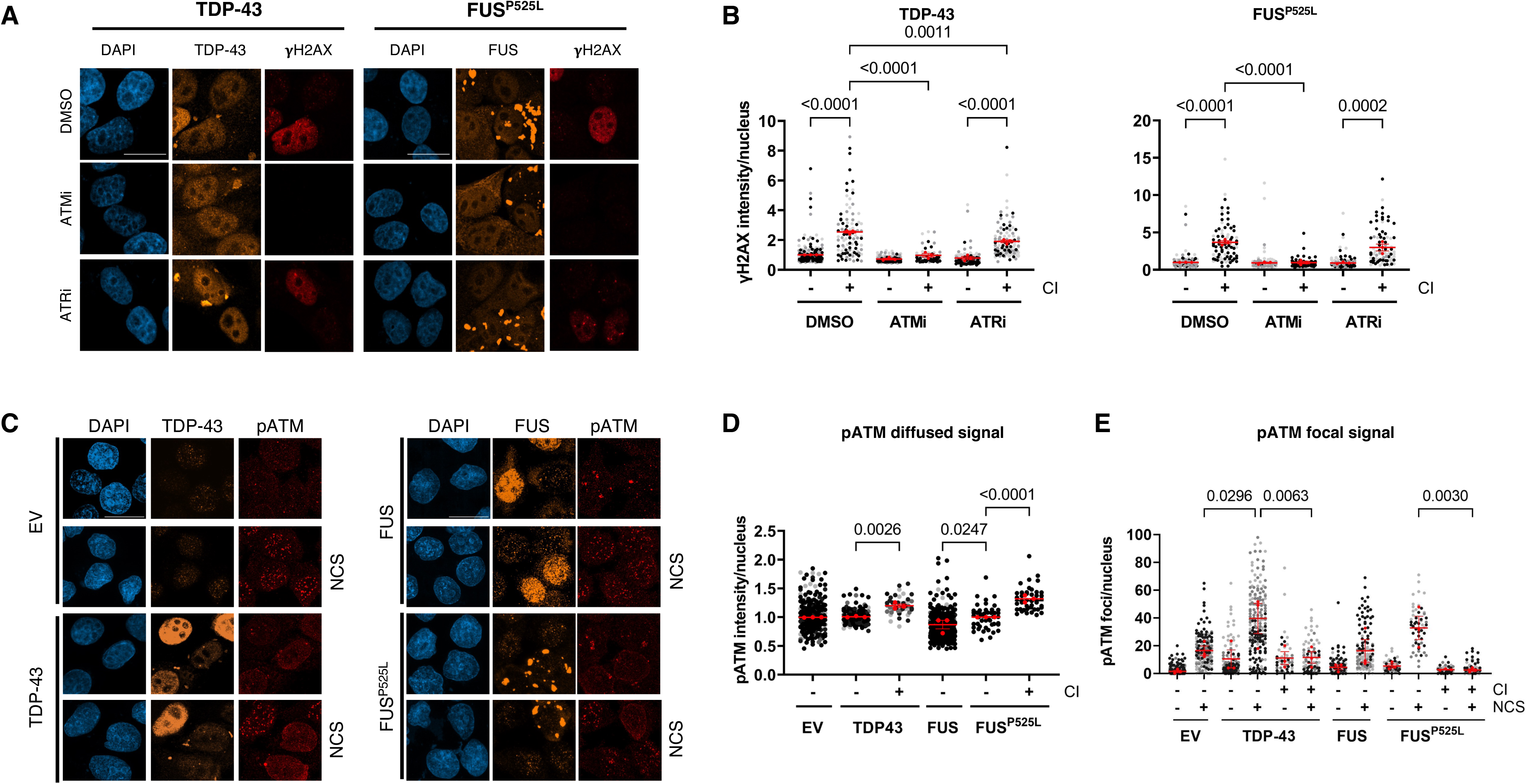
ATM is aberrantly activated, and its inhibition abolishes pan-nuclear γH2AX formation in cells with TDP-43 and FUS^P525L^ CIs. **A)** Analysis by immunofluorescence (IF) of γH2AX levels in HeLa cells expressing TDP-43 or FUS^P525L^ and treated with ATMi or ATRi; cells treated with DMSO were used as a control; nuclei were counter-stained with DAPI; scale bar = 10 µm. **B)** Quantification of γH2AX intensity in HeLa cells with or without cytoplasmic inclusions (± CI) described in A. The red dots of the super-plots represent the mean values of each biological replicate; red bars indicate the average ± SEM of three independent experiments. **C)** IF analysis of pATM signal in HeLa cells transfected with plasmids expressing an empty vector (EV), TDP-43 or FUS^P525L^; nuclei were counter-stained with DAPI; scale bar = 10 µm. **D, E)** or without cytoplasmic inclusions (± CI) determined in C. The red dots of the super-plot represent the mean values of each biological replicate; red bars indicate the average ± SEM of three independent experiments.

We also observed that undamaged cells harboring TDP-43 or FUS^P525L^ CIs exhibited an aberrantly diffused pATM signal, in contrast to those expressing TDP-43 or FUS^P525L^ but lacking CIs (Figure 2C,D). This suggests that the formation of such aggregates is sufficient to trigger a diffused and abnormal ATM activation even without exogenous induction of DNA damage. In line with this, we also observed that ATM activation is inefficient in cells with CIs: while cells without FUS^P525L^ or TDP-43 Cis properly mounted pATM foci formation upon DSB induction following NCS treatment, cells harboring CIs showed a significantly lower number of pATM foci (Figure 2C,E). When we next probed by IF for the activation by phosphorylation of the transducer kinase CHK2, an ATM substrate^34^, upon NCS treatment, CI-bearing cells displayed reduced pCHK2 signals (Figure S2B,C).

In sum, these results indicate that the accumulation of pan-nuclear γH2AX detected in CI-bearing cells relies on a dysfunctional ATM activation, which is responsible for pan-nuclear γH2AX levels but does not efficiently transmit the DDR signal to downstream effector kinases, such as CHK2.

Previous evidence indicates that pan-nuclear γH2AX marks cells in S-phase upon UV-irradiation^35^ and correlates with DNA replication defects^32^. Intriguingly, TDP-43 was shown to associate with replication forks in normal conditions, contributing to protect cells from replicative stress^36^. Although we observed that ATR, often involved in regulating DDR during DNA replication, was dispensable for γH2AX accumulation (Figure 2A,B), we tested if the γH2AX enrichment observed in cells with FUS^P525L^ or TDP-43 CIs might be elicited by replicative stress. To this end, we initially examined the capacity of CI-bearing cells to *de novo* synthesize DNA, by performing a 5-bromo-2’-deoxyuridine (BrdU) incorporation assay followed by IF. We observed that cells with FUS^P525L^ or TDP-43 CIs incorporated much less BrdU, compared to those without CIs or with samples expressing WT FUS or the EV (Figure S2D,E). Similarly, we observed that all the cells harboring FUS^P525L^- or TDP-43-positive CIs had very low or null levels of Cyclin A (Figure S2F), which is expressed in the S/G2-phase of the cell cycle^37,38^, suggesting that such cells were preferentially in G1-phase and not replicating.

These results indicate that CI formation causes DNA damage accumulation and an aberrant ATM activation, which in turn may prevent cells from initiating DNA synthesis and entering the S-phase of the cell cycle.

### FUS^P525L^ and TDP-43 CIs hamper the phosphorylation and the recruitment of 53BP1 at DSBs

53BP1 is one of the main mediators of the DDR cascade and it is involved in DNA repair by non-homologous end-joining (NHEJ), which is the pathway used by non-replicating cells, including neurons, to repair DSBs^39^. Since we previously observed that DDR initiation, as detected by probing for the auto-phosphorylation of ATM, was altered in cells with both FUS^P525L^ and TDP-43 CIs (Figure 2C-E), we tested if the recruitment of 53BP1 was also defective in such cells. While NCS-treated cells without CIs mounted 53BP1 foci, cells with FUS^P525L^ or TDP-43 CIs were apparently unable to form 53BP1 foci upon NCS treatment (Figure 3A,B).

**Figure 3.**
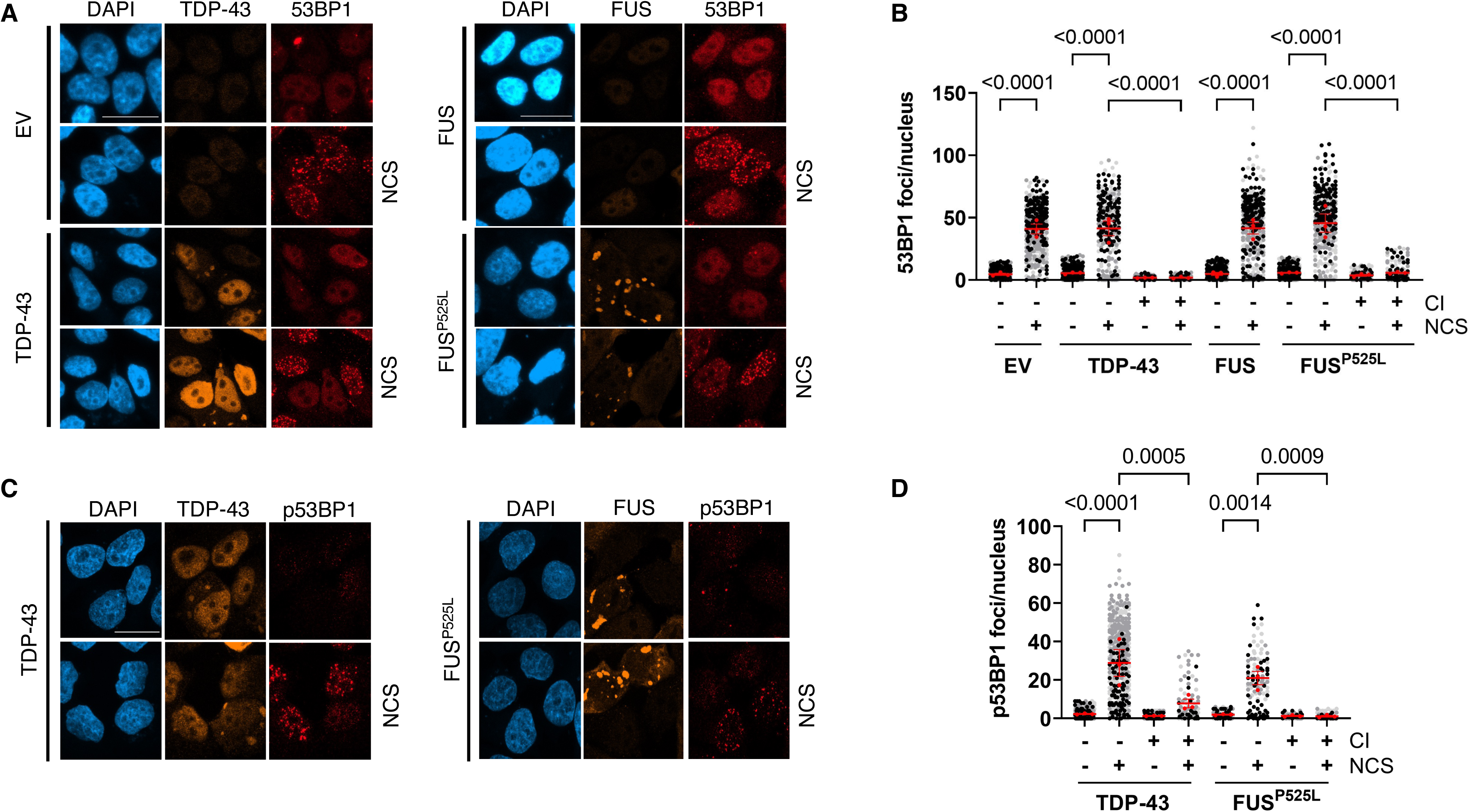
DDR signaling and 53BP1 activation is impaired in cells with TDP-43 and FUS^P525L^ CIs. **A, C)** Representative images of 53BP1 and p53BP1 in damaged (NCS) or undamaged HeLa cells transfected with plasmids encoding for TDP-43, FUS or FUS^P525L^; cells transfected with an empty vector (EV) were used as a control; nuclei were counter-stained with DAPI; scale bar = 20 µm. **B, D)** Quantification of DDR marker signals in cells with or without cytoplasmic inclusions (± CI) determined in A and C, respectively. The red dots of the super-plots shown in B and D represent the mean values of each biological replicate; red bars indicate the average ± SEM of three independent experiments.

Following DSB formation, 53BP1 is phosphorylated by ATM at multiple S/T-Q motifs to modulate DDR and repair^40,41^. We therefore monitored by IF 53BP1 phosphorylation in NCS-treated FUS^P525L^- or TDP-43-expressing cells. We observed that CI-containing cells showed significantly less p53BP1 foci compared to those without aggregates (Figure 3C,D). Differently from what we observed for pATM (Figure 2C,D), undamaged CI-bearing cells showed low or no p53BP1 signal (Figure 3C,D). Also, TDP-43 and FUS^P525L^ aggregation did not substantially affect 53BP1 total levels (Figure S3A), indicating that CIs negatively impacted on 53BP1 activation, rather than on its expression. Overall, these results suggest that the hyper-activation of ATM caused by CI formation is unable to transduce the signal to the downstream DDR mediators.

The scaffold protein MDC1 plays a key role in the DSB signaling, ensuring the recruitment of the E3-ubiquitin ligases RNF8 and RNF168, which in turn allows the assembly of 53BP1 at DSBs^42^. Specifically, MDC1 associates to the site of damage mostly in a γH2AX-dependent manner^43^ and, differently from 53BP1, functions upstream to both homologous recombination (HR), a repair pathway exploited by replicating cells^44^ and NHEJ. To probe for the impact of CI formation on the ability of MDC1 to localize at DSBs, we stained TDP-43- and FUS^P525L^-expressing cells and their controls for MDC1. Differently from 53BP1, NCS-treated cells with CIs showed unaltered MDC1 focal signals (Figure S3B,C).

Taken together, these results indicate that the two DDR mediator proteins, 53BP1 and MDC1, behave differently in cells bearing mutant FUS or TDP-43 CIs, with the profound defect in 53BP1 foci formation not being accompanied by reduced MDC1 focal recruitment.

### Transcription is repressed in cells with TDP-43 or FUS^P525L^ CIs

TDP-43 and FUS functions have been linked to transcription modulation^45^. Moreover, ongoing transcription is halted in response to DSBs, to allow proper DNA damage resolution and preserve genome integrity^46–49^. Specifically, ATM has been shown to initiate transcriptional repression in response to DSB formation^46^. We thus wondered if the increased levels of DNA damage observed in cells with TDP-43 and FUS CIs also correlated with transcriptional inhibition. To monitor *de novo* transcription in cells with CIs, we labelled nascent RNAs in TDP-43- and FUS^P525L^-expressing cell cultures with 5-Ethynyl-Uridine (EU) using a “click-chemistry” approach followed by fluorescent microscopy analysis. We observed a significantly lower EU incorporation signal in cells with either TDP-43 or FUS^P525L^ CIs, indicating a global transcriptional repression (Figure 4A,B).

**Figure 4.**
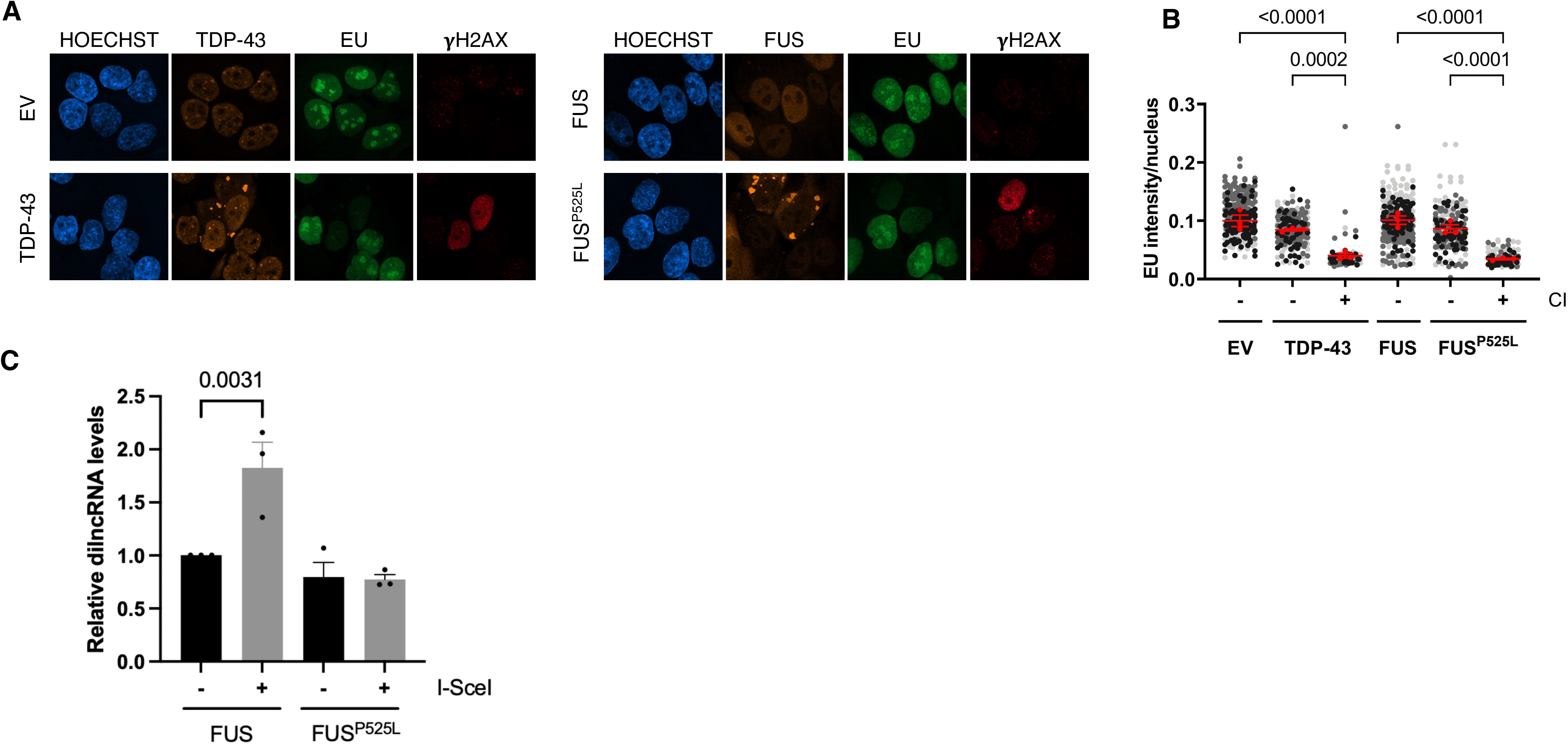
Transcription is halted in cells with TDP-43 and FUS^P525L^ CIs. **A)** HeLa cells expressing TDP-43, FUS or FUS^P525L^ were pulse-labelled with EU and analyzed by immunofluorescence; nuclear DNA was visualized using Hoechst dye. **B)** Quantification of EU signal in cells with or without cytoplasmic inclusions (± CI) shown in A. The red dots of the super-plot represent the mean values of each biological replicate; red bars indicate the average ± SEM of three independent experiments. **C)** dilncRNA expression was studied by strand-specific RT-qPCR in cut (I-SceI +) or uncut (I-SceI -) I-HeLa111 cells selected for the presence of FUS or FUS^P525L^ CIs through FACS. The histogram shows dilncRNA levels normalized over RPLP0 mRNA and shown as relative to the uncut FUS-expressing sample; values are the means ± SEM of three independent experiments.

Since our results indicate that ATM inactivation was sufficient to inhibit γH2AX accumulation in CI-containing cells (Figure 2A,B), we tested its potential impact on global transcription in cells with CIs. We therefore pharmacologically inhibited the ATM kinase activity in HeLa cells expressing TDP-43 or FUS^P525L^ and monitored nascent RNA synthesis using the assay described above. We observed that ATM inhibition elevated the EU signal in cells with CIs, compared to those treated with the vehicle only (Figure S4A,B).

These results therefore indicate that the accumulation of CIs causes an ATM-dependent repression of global transcription.

### Biogenesis of damage-induced non-coding RNAs is defective in CI-containing cells

Following DSB formation, while pre-existing transcription is repressed, RNAPII is recruited at sites of DNA damage where it drives the local synthesis of damage-induced long non-coding RNAs (dilncRNAs)^33,50^. Such transcripts can be further processed by DROSHA and DICER to generate DNA damage response RNAs (DDRNAs) that, along with dilncRNAs, control DDR activation and DNA repair^33,50–52^. We thus investigated if the transcriptional repression observed in CI-containing cells (Figure 4A,B) also impacted on dilncRNA biogenesis. To accomplish this, we profiled the expression of dilncRNAs in I-HeLa111 cells, where a traceable DSB can be generated at a genomic I-SceI recognition site by expressing the I-SceI meganuclease^53^. To enrich and selectively study dilncRNA levels in cells bearing CIs, we took advantage of the Bimolecular Fluorescence Complementation (BiFC) approach^54^. Specifically, we transfected I-HeLa111 cells with two constructs expressing FUS^P525L^ fused to two distinct fragments of the GFP fluorescent protein, mVenus and mCerulean, that emit a fluorescent signal only when brought into proximity with each other upon FUS protein dimer formation, and even more when FUS aggregates^54^ (Figure S4C). Highly fluorescent cells were in this way enriched by fluorescence-activated cell sorting (FACS) and total RNA was isolated from sorted cells and analyzed for dilncRNA expression by strand-specific RT-qPCR. Upon DSB induction with I-SceI, I-HeLa111 cells expressing WT FUS showed elevated dilncRNA levels, while those expressing FUS^P525L^ failed to induce dilncRNAs following DSB generation (Figure 4C).

As mentioned above, along with DICER, the endoribonuclease DROSHA participates in DDRNA biogenesis, and it has been shown to be involved in DDR and repair^51,55–57^. We noticed that cells containing TDP-43 and FUS^P525L^ CIs displayed reduced DROSHA protein levels, compared to those devoid of aggregates (Figure S4D,E). We thus analyzed the expression of *DROSHA* mRNA in cells with FUS aggregates using the BiFC system and observed a significant reduction of its levels in samples expressing FUS^P525L^ (Figure S4F). Contrarily, DICER protein levels were not significantly altered in cells with inclusions (Figure S4G,H).

Taken together, these results indicate that the formation of cytoplasmic aggregates is accompanied by a defective biogenesis of damage-induced non-coding RNAs, important modulators of DDR activation.

### Enoxacin restores proficient DDR, reduces DNA damage accumulation, and partially restores transcription in CI-bearing cells

Enoxacin treatment has been shown to promote the biogenesis of DICER-dependent small RNAs^58^ and we previously demonstrated it stimulates DDR signaling and DNA repair in cultured cells^52,59–61^. Therefore, we tested if enoxacin administration could restore proper DDR activation and reduce DNA damage accumulation in cells with CIs. HeLa cells were treated with enoxacin for 48 hours prior to transfection with TDP-43 or FUS^P525L^ expressing plasmids and monitored for 53BP1 and γH2AX signals by IF. Enoxacin-treated cells bearing TDP-43 and FUS^P525L^ CIs showed both increased 53BP1 foci and reduced γH2AX levels, compared to untreated samples (Figure 5A-D). In addition, supplementing enoxacin to cells expressing TDP-43 decreased DSBs back to normal levels, as revealed by comet assay (Figure S5A,B). These results therefore demonstrate that enoxacin can restore DDR functioning and reduce DNA damage accumulation in cells with CIs.

**Figure 5.**
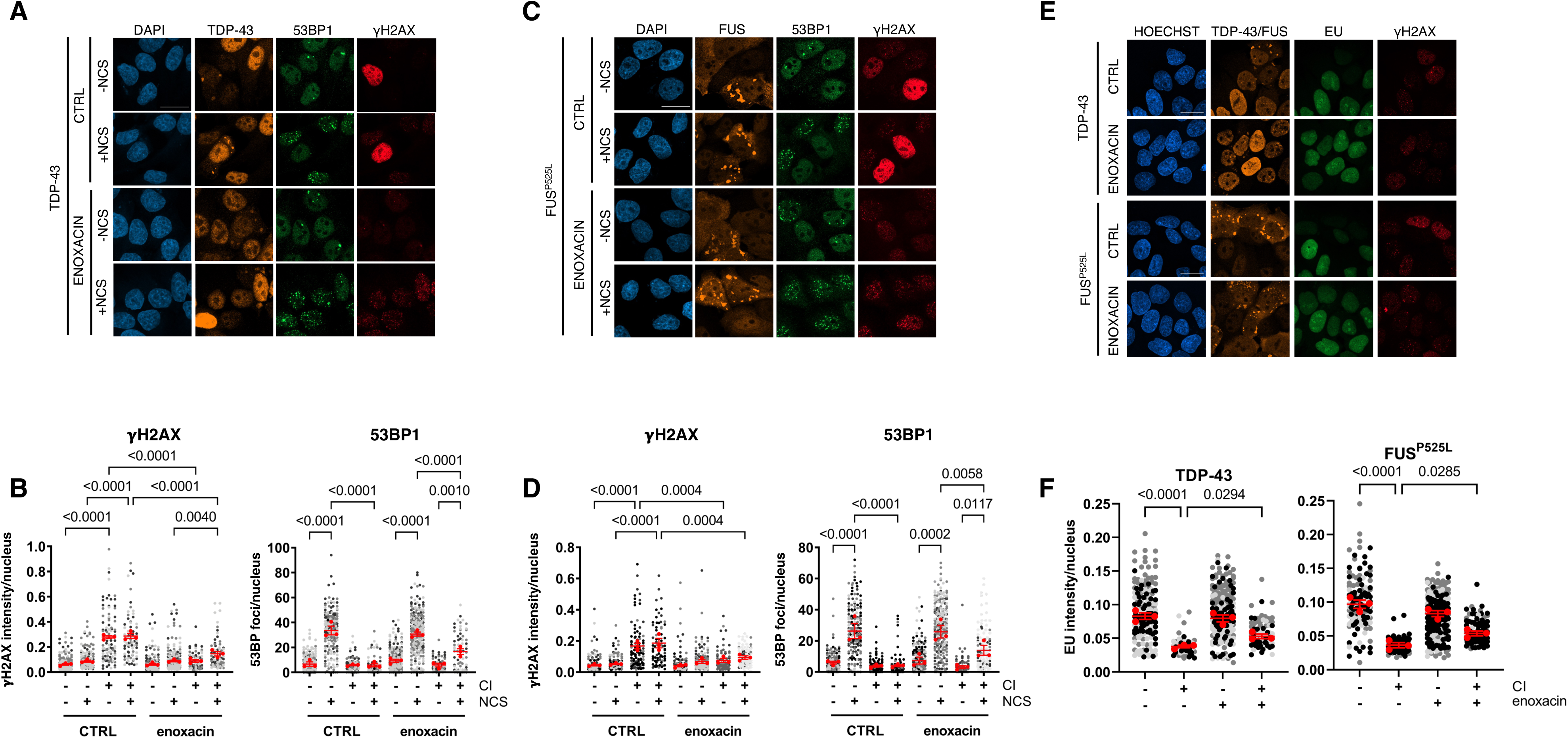
Enoxacin rescues foci formation and partially restores transcription in cells with TDP-43 and FUS^P525L^ CIs. **A, C)** HeLa cells expressing TDP-43 or FUS^P525L^ were treated or not with enoxacin prior to NCS administration and analyzed by immunofluorescence for DDR activation; nuclei were counter-stained with DAPI; scale bar = 10 µm. **B, D)** Quantification of γH2AX intensity and 53BP1 foci in cells with or without cytoplasmic inclusions (± CI) shown in A and C, respectively. The red dots of the super-plots represent the mean values of each biological replicate; red bars indicate the average ± SEM of three independent experiments. **E)** HeLa cells expressing TDP-43 or FUS^P525L^ were treated or not with enoxacin and pulse-labelled with EU prior to immunofluorescence analysis; nuclear DNA was visualized using Hoechst dye; scale bar = 10 µm. **F)** Quantification of EU signal in cells with or without cytoplasmic inclusions (± CI) determined in E. The red dots of the super-plots represent the mean values of each biological replicate; red bars indicate the average ± SEM of three independent experiments.

Since transcription was inhibited in CI-containing cells (Figure 4), we tested whether enoxacin administration could also restore proficient transcription in these cells. Thus, we performed the EU incorporation assay described above in enoxacin-treated cells expressing either TDP-43 or FUS^P525L^ and we observed that enoxacin treatment significantly increased the levels of EU incorporation in cells harboring CIs, with respect to untreated CI-bearing cells, although not to the level detected in cells devoid of CIs (Figure 5E,F).

Altogether, these results show that enoxacin improves DDR and DNA repair in cells with TDP-43 or FUS^P525L^ aggregates, leading to restored global transcription.

### Enoxacin stimulates DDR and lowers DNA damage accumulation in an ALS murine model of FUS pathology

We then sought to extend and validate the observations obtained in cultured cells in *in vivo* models of ALS pathology. To this aim, we administrated enoxacin to either non-pathological heterozygous or pathological homozygous mice expressing the human homolog of FUS (hFUS)^62^ and probed for DDR activation and damage accumulation, by staining murine spinal cords for γH2AX and 53BP1 by immunohistochemistry (IHC) techniques. Consistent with our results in cultured cells, hFUS homozygous mice presented higher levels of γH2AX and reduced 53BP1 signals in their spinal cord, with respect to heterozygous animals (Figure 6A,B). Importantly, in enoxacin-treated homozygous mice we observed that the signal of 53BP1 increased, whereas γH2AX levels decreased, compared to vehicle-treated animals (Figure 6A,B).

**Figure 6.**
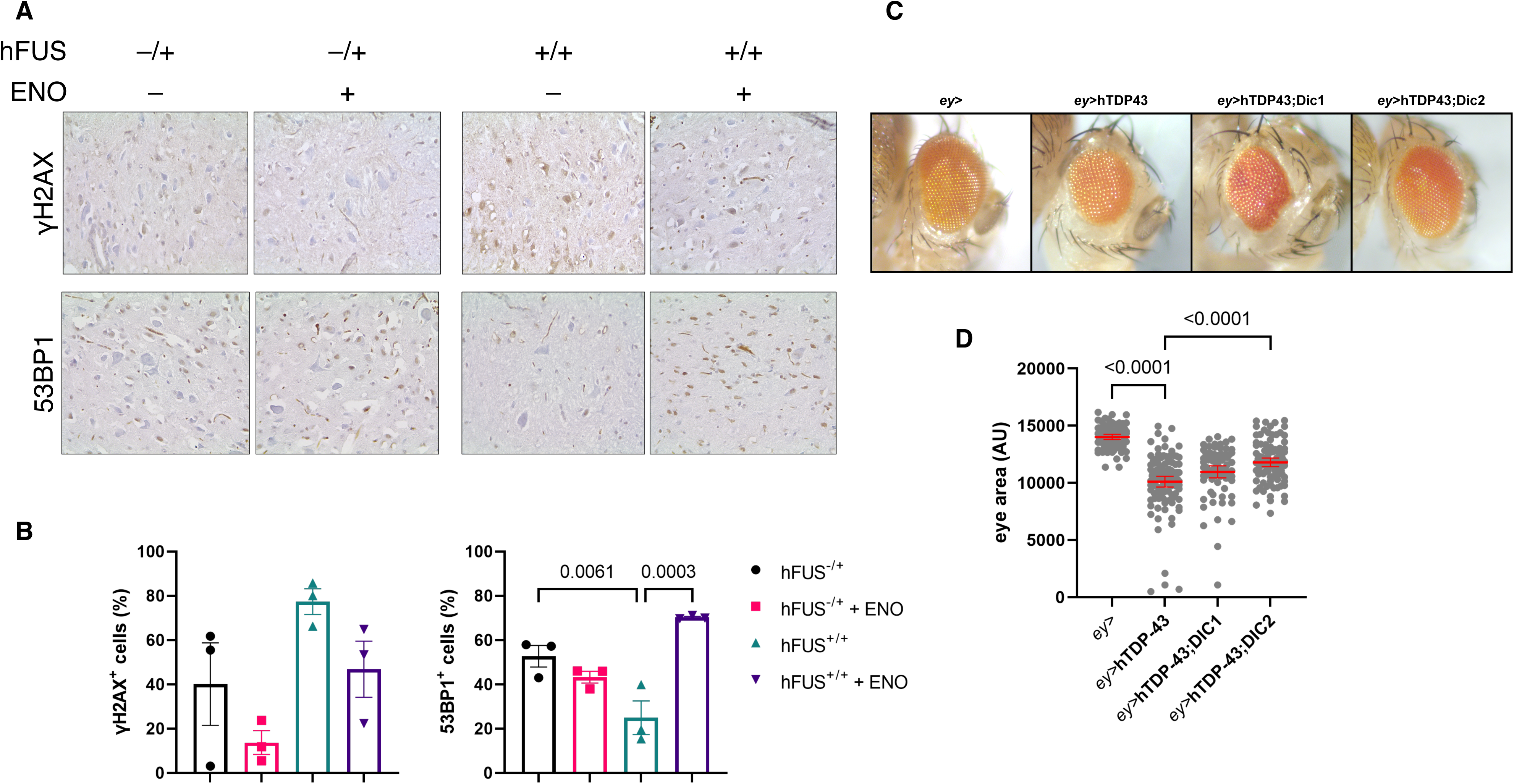
Enoxacin stimulates DDR and reduces DNA damage accumulation in hFUS murine spinal cords. Dicer-2 overexpression counteracts retina degeneration in hTDP-43 *Drosophila melanogaster* eyes. **A)** Representative IHC images of spinal cords of hFUS-expressing mice, treated with enoxacin or vehicle and stained for γH2AX and 53BP1. **B)** Quantification of the percentage of cells positive for the indicated markers as determined in A. Values are the means ± SEM; three mice were studied for each condition. **C)** Representative images of *Drosophila melanogaster* eyes from adult flies expressing the indicated constructs under control of eyeless-GAL4.Dot plot showing the quantification of mean eye area of the flies with the indicated genotypes; more than 70 eyes per genotype were analyzed in three independent experiments; red bars represent the means ± SEM.

Taken together, these exciting results suggest that enoxacin administration *in vivo* in mice with FUS pathology was effective at reducing DNA damage accumulation.

### DICER overexpression counteracts neurodegeneration in a *Drosophila melanogaster* ALS model system

Our results in mice show that the enhancement of DNA repair through enoxacin administration reduces DNA damage accumulation *in vivo*. However, how this can impact on neurodegeneration remains unexplored. To test this, we used a *Drosophila melanogaster* model of TDP-43-mediated neurotoxicity, in which the ectopic expression of the human TDP-43 (hTDP-43) in the fly ommatidia causes progressive retinal degeneration, as shown by a significant reduction of the eye area compared to control flies^63^ (Figure 6C,D). To mimic the effects of enoxacin administration in this system, we overexpressed both *D. melanogaster* isoforms of Dicer in hTDP-43-expressing flies and monitored their impact on retinal degeneration. Differently from mammals, the *D. melanogaster* genome encodes for two distinct Dicer enzymes: Dicer-1, mainly involved in cytoplasmic microRNA (miRNA) biogenesis, and Dicer-2 with a more prominent role in short interfering RNA (siRNA) processing^64^. Remarkably, the overexpression of Dicer-2 significantly rescued the eye area in hTDP-43-expressing flies (Figure 6C,D). Conversely, no major effects were observed in flies co-expressing hTDP-43 and Dicer-1, ruling out a central role for miRNAs in preventing retinal degeneration (Figure 6C,D).

These results indicate that stimulating Dicer-2 functions may be beneficial in counteracting TDP-43-mediated neurodegeneration.

## DISCUSSION

Accumulation of unrepaired DNA damage and the resulting disruption of genome integrity can have profound consequences, including ageing and inflammation, which in turn increase the susceptibility to various pathological conditions, especially neurodegenerative diseases^23,65,66^. In light of this, mounting evidence have recently put into relation DNA damage repair defects with neurodegeneration in ALS^23^. ALS is characterized by an aberrant accumulation of insoluble cytoplasmic inclusions (CIs) containing TDP-43 and other RNA-binding proteins, including FUS^2^. Interestingly, FUS and TDP-43 have been reported to play crucial roles in DNA damage repair^67^: FUS associates with DNA lesions by interacting with PAR-chains generated at damaged chromatin^68^, and favors DNA damage resolution by recruiting the XRCC1/Ligase lII repair complex to DNA lesions^69^. In addition, recently, TDP-43 has been implicated in DNA repair and reported to associate at DSBs with factors of the non-homologous end-joining (NHEJ) machinery, ultimately contributing to DNA damage resolution^70^. These studies have highlighted a causative link between the depletion of FUS and TDP-43 with genome instability and reduced survival in neuronal cells^69,70^. Nonetheless, the impact of TDP-43 and FUS cytoplasmic aggregation – a pathological hallmark of ALS – on DNA damage accumulation has not been fully elucidated.

Here, we show that the deposition of TDP-43 and FUS^P525L^ CIs, which are associated with stress granules (SGs) (Figure S1E), triggers a dysfunctional DDR (Figure 2C-E and 3), and induces DSB accumulation (Figure 1C-G). Differently, cells overexpressing FUS or TDP-43, but devoid of CIs, do not accumulate DNA damage (Figure 1C-G) and display proficient DDR when exogenous DNA damage is inflicted (Figure 2C-E and 3). Specifically, cells accumulating CIs display a hyper-activation of the apical DDR kinase ATM, responsible for the massive increase of the pan-nuclear γH2AX signal observed (Figure 2). Such aberrant activation of ATM is however unable to transduce the signal to the downstream DDR and repair factors, including 53BP1 (Figure 3), ultimately hampering their recruitment to DNA damage site and consequently DNA damage resolution through NHEJ, the predominant DNA repair pathway in non-dividing cells. Importantly, ATM inhibition using small molecules prevents the accumulation of DNA damage signals in cells with CIs (Figure 2A,B). Our results therefore put in a causative relationship the observed genotoxicity and TDP-43 and FUS cytoplasmic aggregation.

Shutdown of global transcription is one of the immediate consequences of DNA damage generation^71^. In particular, the activation by auto-phosphorylation of ATM in response to DNA damage has been shown to enforce chromatin modifications surrounding DSBs, which in turn prevent RNAPII elongation^46^. In addition, loss of either TDP-43 and FUS has been linked to transcription arrest and consequent formation of DNA damage at sites of active transcription^45^. Consistent with this, cells harboring TDP-43 or FUS^P525L^ CIs display dysfunctional ATM activation (Figure 2C-E) and pan-nuclear γH2AX signals (Figure 1C-E, 2A,B and 4A) and exhibit defective global transcription (Figure 4A,B). Pharmacological inactivation of ATM rescues proficient RNA synthesis in CI-containing cells (Figure S4A,B). Overall, our results indicate that TDP-43 or FUS^P525L^ CI formation, by inducing aberrant DDR and DSB generation, hampers transcription. Notably, the observed CI-induced transcriptional repression may have profound implications particularly in neurons, where longer transcripts are enriched with respect to other cell types^72,73^. In this regard, the inactivation of genes involved in regulating RNAPII processivity, by favoring the accumulation of truncated transcripts, has been associated to synaptic dysfunction, autism, and neurodegeneration^73^. Such findings, along with the results shown here, provide an intriguing explanation for the higher sensitivity of neural tissues to CI accumulation, compared to others.

Interestingly, the formation of cytosolic aggregates not only hampers the synthesis of damage-induced non-coding RNAs (Figure 4C), but it might also impair their processing by reducing the expression of DROSHA (Figure S4D-F). Such events may ultimately interfere with faithful DNA damage resolution^33,50,57,74^, likely establishing a detrimental feedback loop that further exacerbates the accumulation of unrepaired DNA lesions. Accordingly, in this double-hit process, even minimal improvements in DDRNA biogenesis may exert beneficial effects.

The small molecule enoxacin, which was reported to enhance DICER activity^58^, is one of the few compounds able to boost DDR and improve repair^52,59–61^. Its administration was demonstrated effective in improving DNA damage resolution in irradiated primary mouse cortical neurons^52^ and in an AD model of human neuronal cells^60^. In the present study, we show that stimulating DICER activity through enoxacin restored functional DDR and decreased DNA damage accumulation in HeLa cells with TDP-43 and FUS^P525L^ CIs, and *in vivo* in hFUS-expressing mice (Figure 5A-D and 6A,B). To evaluate whether the observed effects of DICER enhancement on stimulating DDR and repair were also beneficial at counteracting neurotoxicity, we expressed both Dicer-1 and Dicer-2 in a *Drosophila melanogaster* model of TDP-43-mediated retinal neurodegeneration. Intriguingly, while the overexpression of Dicer-2 almost completely restored the normal fly eye size, Dicer-1 ectopic expression had no impact (Figure 6C,D). This suggests that the small RNAs generated by DICER exert their function in controlling DDR and repair, ultimately promoting neuronal cell survival through an siRNA-mediated mechanism, rather than in a miRNA-like fashion.

In conclusion, our results strongly suggest that the factors involved in DDR may represent relevant pharmacological targets for the treatment of the cytotoxic effects caused by TDP-43 and FUS CIs.

## ACKNOWLEDGEMENTS

We would like to thank Dr Simone Sabbioneda (IGM-CNR, Pavia) for support with the fluorescence-activated cell sorting. **S.Fa** is supported by Fondazione Umberto Veronesi. **S.M.** is supported by Istituto Universitario di Studi Superiori (IUSS, Pavia, Itay). **N.D’A.** is supported by: Fondazione AriSLA ETS (Fondazione di Ricerca per la SLA ETS), SwitchALS project; #NEXTGENERATIONEU (NGEU) and founded by the Ministry of University and Research (MUR), National Recovery and Resilience Plan (NRRP), project MNESYS (PE0000006) – A Multiscale Integrated Approach to the Study of the Nervous System in Health and Disease (DN. 1553 11.10.2022). **M.C.** is supported by: Fondazione AriSLA ETS (Fondazione di ricerca per la SLA ETS), SwitchALS project; CNR (DBA.AD005.225-NUTRAGE-FOE2021). **G.Ce.** is supported by: CNR (DBA.AD005.225-NUTRAGE-FOE2021); CNR (Flagship Project Interomics); EU funding within the MUR PNRR “National Center for Gene Therapy and Drugs based on RNA Technology” (# CN00000041 CN3 RNA). **F.d’A.d.F.** is supported by: ERC advanced grant (TELORNAGING—835103); AIRC-IG(21762); Telethon (GGP17111); AIRC 5×1000 (21091); Progetti di Ricerca di Interesse Nazionale (PRIN) 2015“ATR and ATM-mediated control of chromosome integrity and cell plasticity”*(2015SJLMB); Progetti di Ricerca di Interesse Nazionale (PRIN) 2017 “RNA and genome Instability” (2017NWEXEP); Progetto AriSLA 2016 “DDRNA&ALS and 2021 “DDR&ALS” (FG_24/2020); POR FESR 2014-2020 Regione Lombardia(InterSLA project) (DSB.AD004.294); FRRB—Fondazione Regionale per la Ricerca Biomedica—under the frame of EJP RD, the European Joint Programme on Rare Diseases with funding from the European Union’s Horizon 2020 research and innovation program under the EJP RD COFUND-EJP NO 825575 (EJPRD19-206); co-funding European Union – Next Generation EU, in the context of The National Recovery and Resilience Plan, Investment Partenariato Esteso PE8 “Conseguenze e sfide dell’invecchiamento”, Project Age-It (Aging Well in an Aging Society) GAE 492 PNRR PE_8 SPOKE 2; PNRR-CN3 “National Center for Gene Therapy and Drugs based on RNA Technology”; ERC POC TELOVACCINE – 101113229; Telethon GMR23T2007. **U.G.** is supported by: Progetto AriSLA 2016 “DDRNA&ALS and 2021 “DDR & ALS”; PRIN 2020 CXFL4T; Istituto Superiore di Sanità RIPREI2023_7c8ae10d783c. **S.Fr.** is supported by: Progetto AriSLA 2016 “DDRNA&ALS” and 2021 “DDR&ALS”; PNRR-CN3 “National Center for Gene Therapy and Drugs based on RNA Technology”; POR FESR 2014-2020 Regione Lombardia (InterSLA project).

## AUTHOR CONTRIBUTIONS

**S.Fa.** carried out immunofluorescence analysis for quantification of WT and mutant FUS CIs; experiments of colocalization between FUS and stress granules; immunofluorescence analysis of γH2AX signals with and without ATM and ATR inhibitors, analysis of pATM, 53BP1 and p53BP1 signals in cells expressing WT and mutant FUS and immunoblot experiments for FUS expression upon transfection in HeLa cells. **F.E.** performed immunofluorescence analysis for γH2AX; ATM and ATR inhibition experiments; BrdU and CycA staining; EU incorporation assays; immunofluorescence and RT-qPCR analysis for DROSHA expression; enoxacin treatment experiments and generated the constructs for the BiFC assays. **S.M.** performed the immunoblot analyses to quantify TDP-43 and FUS levels, carried out quantitative immunofluorescence analyses for pATM, pCHK2, 53BP1, p53BP1, MDC1, DROSHA and DICER in HeLa cells; analyzed dilncRNA expression and edited the manuscript. **O.B.** carried out the experiments concerning the impact of mutant TDP-43 isoforms on cytoplasmic inclusion and stress granule formation. **M.D.S.** carried out the experiments involving *Drosophila melanogaster* strains. **I.D.V.** conducted the experiments in mice and generated the spinal cord tissues from such animals. **G.Ci.** performed IHC analyses on murine specimens. **A.R.** carried out the EU incorporation assays with ATM inhibitors. **F.P.** conducted the IHC staining in murine samples. **A.G.** assisted S.M. with the fluorescence-activated cell sorting. **N.D’A.** and **M.C.** coordinated the experiments in hFUS mice, supervised I.D.V. and edited the manuscript. **G.Ce.** coordinated the analyses in *Drosophila melanogaster* strains, supervised M.D.S. and edited the manuscript. **F.d’A.d.F.** participated in designing the study and edited the manuscript. **U.G.** conceived, conducted and analyzed the comet assays; contributed to the study design, assembled and revised the manuscript. **S.Fr.** conceived the study, coordinated experimental activities, wrote and edited the manuscript.

## COMPETING INTERESTS

The authors declare no competing interests.

## MATERIALS AND METHODS

### Cell culture and treatments

HeLa cells were cultured in DMEM supplemented with 10% FBS, 1% L-glutamine and 1% penicillin/streptomycin (P/S). DNA damage was induced by treating cells with 50 ng/mL neocarzinostatin (NCS) (Sigma, #N9162) for 20 minutes at 37 °C. I-HeLa111 were cultured in DMEM supplemented with 10% tetracycline-free FBS, 1% L-glutamine and 1% P/S and selected regularly with G418 (1 mg/ml) and hygromycin B (300 μg/ml). In I-HeLa111, I-SceI-mediated double-strand breaks were generated by doxycycline administration (1 μg/ml) for at least 16 h. To inhibit ATM and ATR, cells were treated with 5 µM KU60019 and 10 µM VE-821 overnight, respectively. Where indicated, HeLa cells were treated with 50 µM enoxacin (Millipore, #557305-250MG) for 24 h prior to plasmid transfection and left until the end of the experiment (48 h in total).

### Transfections

1 μg of each plasmid was transfected for 24 hours using Lipofectamine 2000 (Invitrogen; #11668-027) according to the manufacturer instructions. The following plasmids were employed: pcDNA3.1+ (ThermoFisher) was used as an empty vector negative control; MYC-tagged WT TDP-43 expressing vector, a kind gift of Dr. Emanuele Buratti on concession of Dr. Leonard Petrucelli (Mayo Clinic, Jacksonville, Florida); FLAG-tagged WT FUS and FUS^P525L^ expressing plasmids. For the Bimolecular Fluorescence Complementation (BiFC) experiments, WT FUS or FUS^P525L^ were cloned in pCS2plus-mCerulean 156-239-GGGS (Addgene, #162616) and pCS2plus-mVenus1-155-GGGS (Addgene, #162610), both gift from Marco Morsch^54^, using the Gibson Assembly (ThermoFisher). 0.7 µg of each vector were transfected using Lipofectamine 2000.

### Immunofluorescence and image analysis

Immunofluorescence for DDR markers was performed as already described^51^. Images were acquired using a confocal linear laser-scanning microscope (Zeiss LSM 800). Quantitative immunofluorescence analyses were performed in parallel with identical acquisition parameters and exposure times among conditions, using CellProfiler software^75^. DICER cytoplasmic signal was quantified using Fiji software (NIH). For a complete list of the antibodies used in this study see Table 1.

**Table 1:**
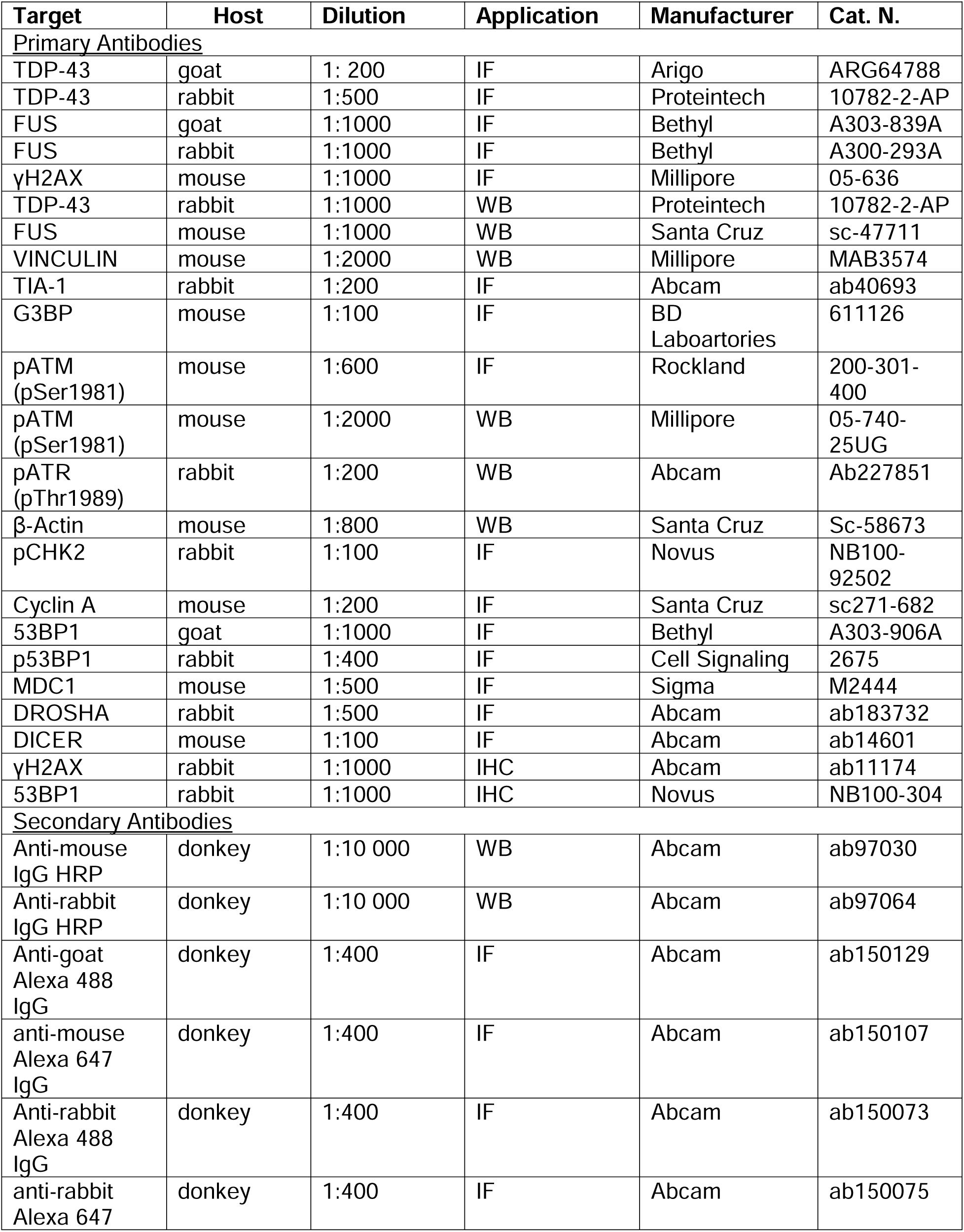
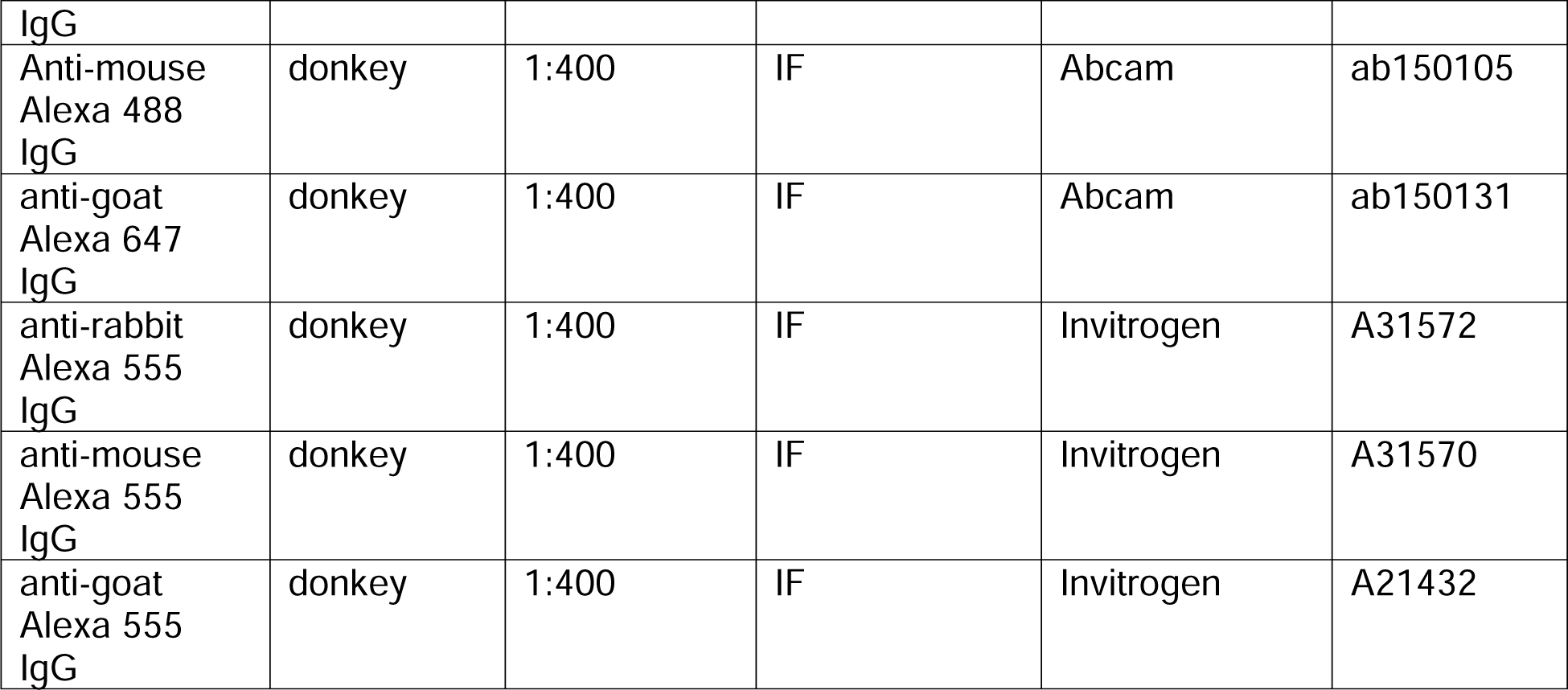
Antibodies used in this study.

### Immunoblotting

Immunoblotting analyses were conducted as shown previously^57^. Protein amounts were quantified using Image Lab 6.1 software (Bio-Rad). For a complete list of the antibodies used in this study see Table 1.

### Comet assay

To detect DSB formation in cultured cells, we performed a neutral comet assay as previously described^52^.

### BrdU incorporation assay

After plasmid transfection, cells were treated with 10 µM BrdU (MedChemExpress, #HY-15910/CS-2028) overnight and incubated at 37°C. The following day, cells were washed twice in 1X PBS and fixed in 4% paraformaldehyde for 10 minutes at RT. Coverslips were then washed twice in 1X PBS and blocked in 1X PBG (0.5% bovine serum albumin, 0.2% gelatin from cold water fish skin (Sigma Aldrich, #G7765) in 1x PBS) for 1h at RT. Next, coverslips were incubated for one hour at RT in a humid chamber with a 1X PBG mix containing an anti-BrdU primary antibody (BD Laboratories, #347580) diluted 1:20 and RQ1 DNAse (Promega, #M610A) and RQ1 DNAse buffer (Promega, #M198A), each diluted 1:10. After incubation of the primary antibody, cells were washed three times in 1X PBS and immunofluorescence continued as described above.

### EU incorporation

5-EU incorporation experiments were performed as already described^76^. Briefly, following DNA damage generation via NCS treatment, nascent RNAs were labelled with 1 hour pulse of 1 mM 5-EU. Cells were then fixed in 4% paraformaldehyde (PFA; ChemCruz; #sc-281692) and levels of EU incorporation were detected by Click-iT RNA imaging kit (Invitrogen; #C10330) according to the manufacturer’s instructions.

### Cell sorting

24 h post plasmid transfection, cells were washed twice with 1X PBS, gently scraped and collected in 1 mL of sorting buffer (2 mM EDTA, 20 mM HEPES pH 7.3 in 1X PBS) per condition, collected in 15 mL tubes and placed on ice. Next, samples were transferred onto a 5 mL falcon equipped with a 35 µM nylon mesh (Corning, #3532235) and vortexed thoroughly to dissociate any cell clusters that might have formed. Next, samples were analysed on an S3e Cell sorter (BioRad). 30000 cells transfected with the EV were analysed first to set the gates. Next, cells expressing WT or FUS^P525L^ were sorted in purity mode and the positive cells were collected. Following cell sorting, cells were centrifuged at 4°C for 1.5 minutes at full speed and the supernatant was discarded. Cells were then either lysed to collect RNA or frozen at –80 °C for future use.

### RT-qPCR

Total RNA extraction was performed using the miRNeasy Mini Kit (Qiagen, #1038703), according to the manufacturer’s instructions. 1 µg of total RNA was reverse transcribed using the SuperScript IV First-Strand Synthesis System (Invitrogen, #18091050). For dilncRNA detection in I-HeLa111 cells, 500 ng of total RNA were reverse transcribed with the SuperScript III First-Strand Synthesis System (Invitrogen, #18080-51) using gene-specific primers. Then, samples were chilled on ice and incubated for 20 minutes at 37°C with 1 µL of RNase H (Invitrogen). Real-time quantitative PCR (qPCR) reactions were performed on a LightCycler® 480 II Sequence Detection System (Roche) using QuantiTect SYBR® Green PCR kit (Qiagen; #204145). For a complete list of primers used in this study see Table 2.

**Table 2:**
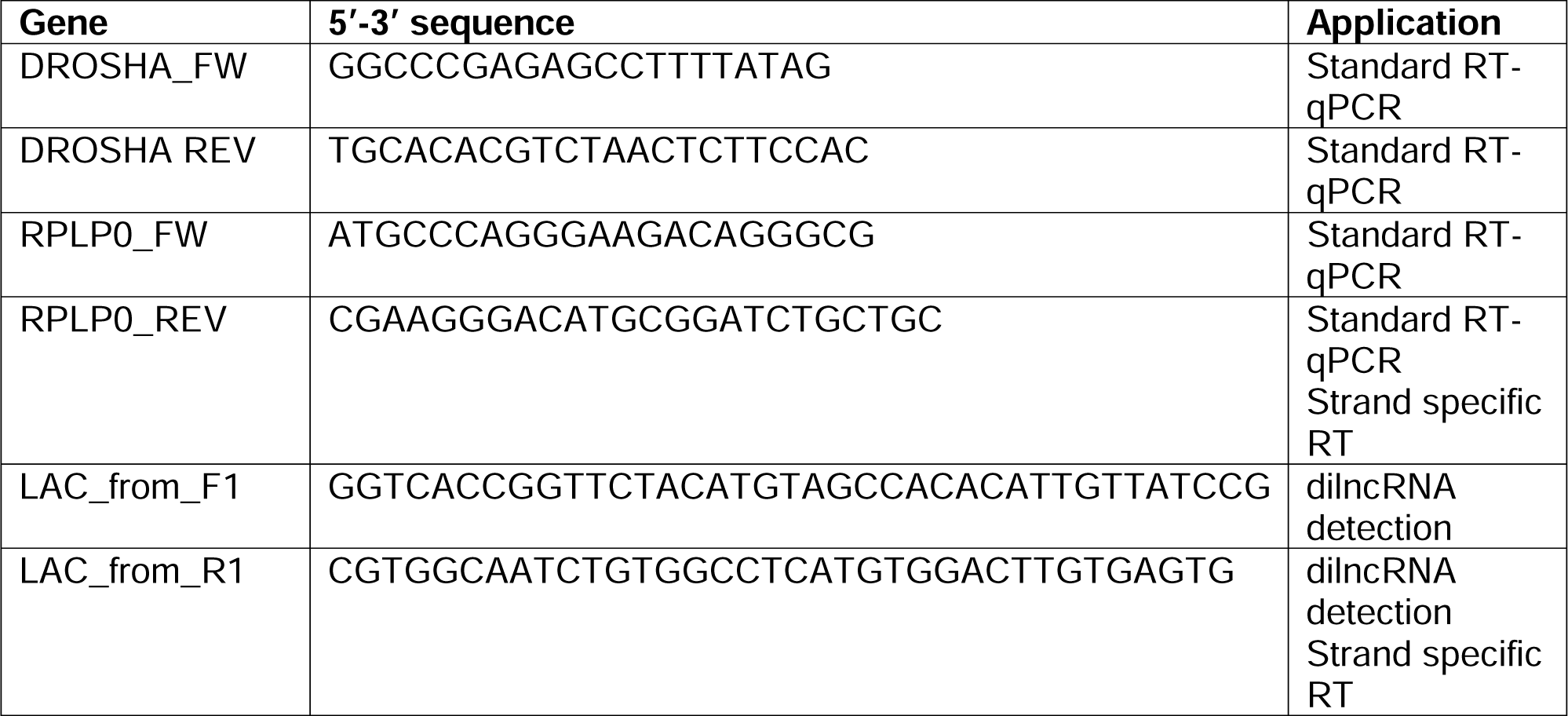
Primers used in this study.

### Mice

Mice were housed at the Tor Vergata University Animal Facility (CIMETA) in accordance with the FELASA Recommendations, European Guidelines for the use of animals in research (2010/63/EU), and Italian Laws (D.L. 26/2014). They were kept at constant temperature of 22 ± 1 °C, relative humidity of 50%, and a 12-h light cycle (7 a.m.–7 p.m.), with free access to food and water. To ensure nutrition and hydration, wet food was provided to cages where mice displayed signs of paralysis. Hemizygous Tg (Prnp-FUS) WT3Cshw/J mice expressing human wild-type FUS (hFUS +/-, Jackson Laboratories), displaying no pathological signs, were backcrossed to obtain homozygous mice (hFUS+/+), models of ALS. Genotyping was performed as previously described^62^. All experiments were conducted in compliance with the ARRIVE guidelines, the European Guidelines (2010/63/EU) and the requirements of Italian laws (D.L. 26/2014). Every effort was made to minimize mice suffering and reduce the number of mice used to obtain reliable results. A total of n = 12 hFUS mice were included in the study. They were treated intraperitoneally daily with either enoxacin at a dose of 10 mg/kg or with vehicle. Enoxacin (Millipore, #557305-250MG) was initially dissolved in 1 M NaOH solution to a concentration of 312 mM and then diluted 100× in PBS for injection. Vehicle was 10 mM NaOH in PBS. Treatment was initiated at about 26 days, which correspond to early symptoms onset in hFUS +/+ mice, for the following 8 days, until mice reached an advanced symptomatic stage (about 34 days). Mice were euthanized with CO_2_, spinal cords were removed and fixed in a 4% paraformaldehyde (PFA) for 12 h and finally immersed in a 30% sucrose solution in PBS.

### Immunohistochemistry

Immunohistochemistry staining was conducted as shown before^77^. For each section studied, five non-overlapping fields were analyzed using the segmentation based algorithm Multiplex IHC v3.2.3 of HALO software (Indica Labs).

### Fly strains

*Drosophila* stocks and crosses were maintained on *Drosophila* standard medium (Nutri-fly, Genesee Scientific) at 25°C. UAS-hTDP-43 fly line was gifted from Fabian Feiguin. Bloomington stock center (http://flybase.bio.indiana.edu/) provided the GAL4 driver and the other strains utilized: eyeless-GAL4 inserted on the second chromosome (5534 (w[*]; P{w[+m*]=GAL4-ey.H}3-8); both Dicer transgenes were inserted on the third chromosome (59022 (w*;P{UAS-Dcr-2.D}2,P{UAS-mCherry.CAAX.S}2;TM2/TM6B,Tb1)), and (36510 (y1w*;P{UAS-Dcr-1.D}3).

### Statistical analysis

Prism 9 software (GraphPad) was used to generate graphs, to perform statistical analysis and to occasionally remove values with the Robust regression and Outlier removal method. Statistical analysis was performed with the One-way ANOVA test, unless differently indicated.

---

**Figure S1 (related to Figure 1).**
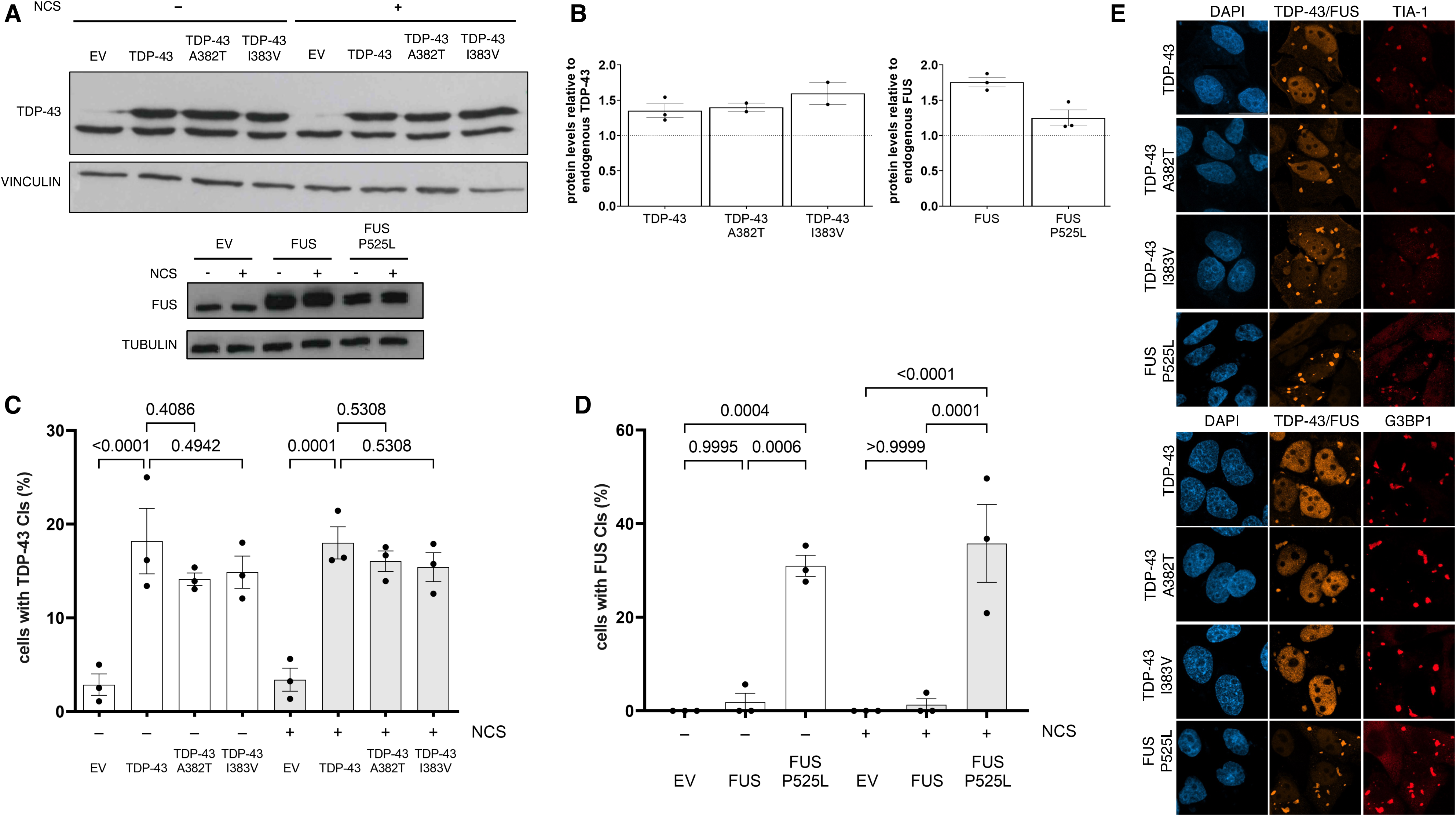
TDP-43 and FUS^P525L^ CIs colocalize with SGs. **A)** Analysis by immunoblot of wild-type or mutant TDP-43 or FUS levels in damaged (NCS +) or undamaged (NCS -) HeLa cells. **B)** The histograms show the ratio between the transfected TDP-43 and FUS protein levels and their endogenous protein amounts as determined in A; values are the means ± SEM of at least 2 independent experiments. **C, D)** Histograms showing the percentage of HeLa cells harboring wild-type or mutant TDP-43 of FUS CIs. **E)** Immunofluorescence images of wild-type, mutant TDP-43, or FUS^P525L^ CIs (orange) co-localizing with stress granules (SGs), labelled by TIA-1 or G3BP1 staining (red); nuclei were counter-stained with DAPI; scale bar = 20 µm.

**Figure S2 (related to Figure 2).**
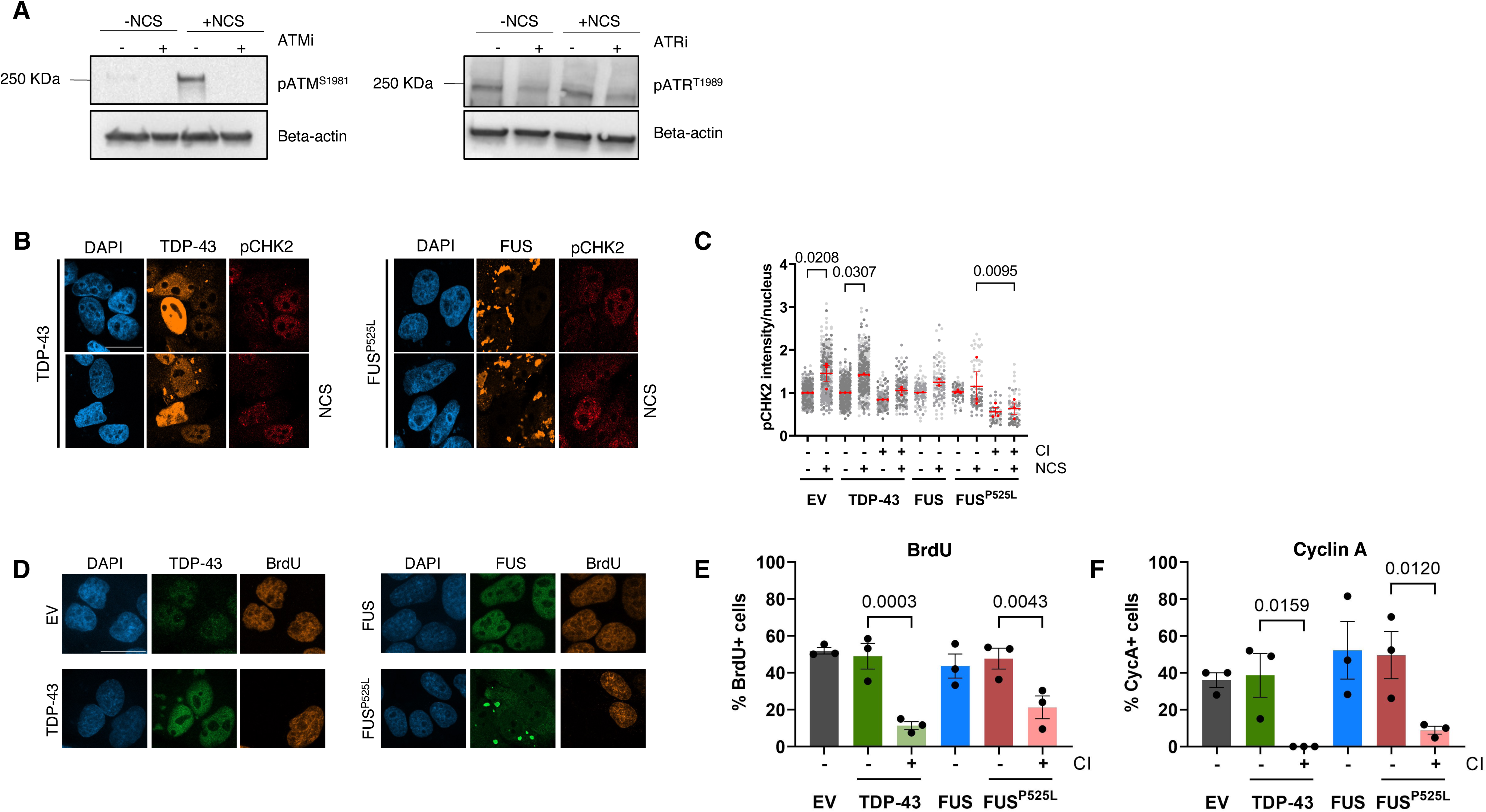
Cells with TDP-43 and FUS^P525L^ CIs arrest preferentially in G1. **A)** Analysis by immunoblot of ATM and ATR activation (pATM^S1981^ and pATR^T1989^, respectively) in damaged (+NCS) or undamaged (-NCS) HeLa cells treated with ATMi or ATRi; cells treated with DMSO only (-) were used as a control. **B)** Representative images of pCHK2 signals in damaged (NCS) or undamaged HeLa cells transfected with plasmids encoding for TDP-43 or FUS^P525L^; nuclei were counter-stained with DAPI; scale bar = 10 µm. **C)** Quantification of pCHK2 signals in cells with or without cytoplasmic inclusions (± CI) determined in B. The red dots of the super-plot represent the mean values of each biological replicate; red bars indicate the average ± SEM of three independent experiments. **D)** Immunofluorescence images of BrdU incorporation assay in HeLa cells transfected with plasmids expressing TDP-43, FUS or FUS^P525L^; cells transfected with an empty vector (EV) were used as a control; nuclei were counter-stained with DAPI; scale bar = 10 µm. **E)** Histograms showing the percentage of BrdU-positive cells with or without cytoplasmic inclusions (± CI) determined in D; values are the means ± SEM of three independent experiments. **F)** The histograms show the percentage of cells, containing or not cytoplasmic inclusions (± CI), that express Cyclin A (CycA+); values are the means ± SEM of three independent experiments.

**Figure S3 (related to Figure 3).**
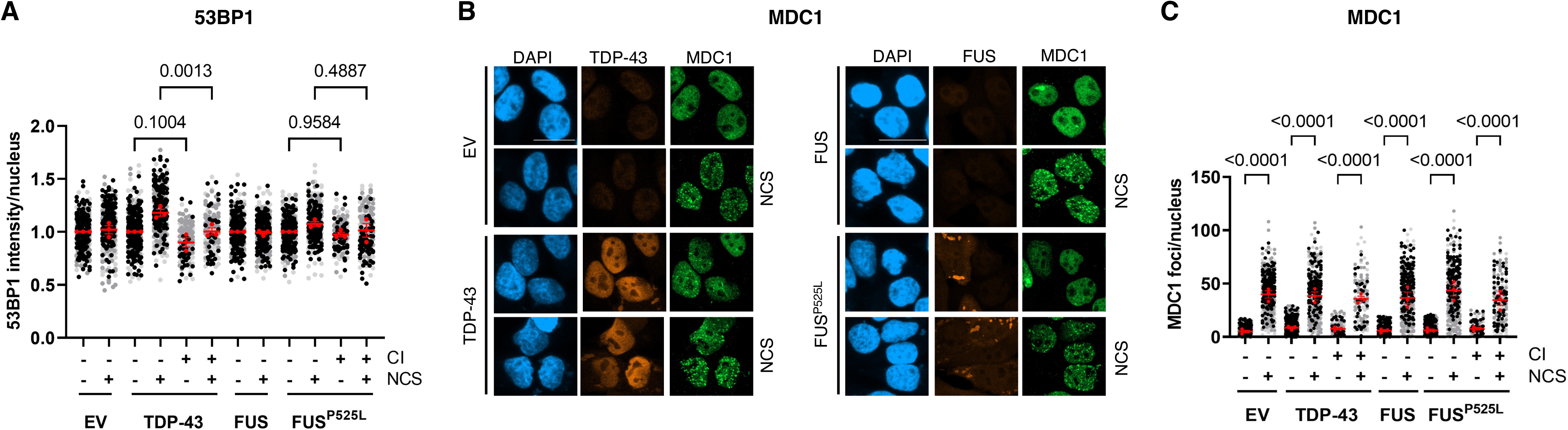
**A)** Quantification of 53BP1 intensity in damaged (NCS +) and undamaged HeLa cells with or without cytoplasmic inclusions (± CI) determined in Figure 3A. The red dots of the super-plot represent the mean values of each biological replicate; red bars indicate the average ± SEM of three independent experiments. **B)** Immunofluorescence analysis of MDC1 recruitment in damaged (NCS) or undamaged HeLa cells transfected with plasmids encoding for TDP-43, FUS or FUS^P525L^; cells transfected with an empty vector (EV) were used as a control; nuclei were counter-stained with DAPI; scale bar = 10 µm. **C)** Quantification of MDC1 foci in cells with or without cytoplasmic inclusions (± CI) determined in B. The red dots of the super-plots represent the mean values of each biological replicate; red bars indicate the average ± SEM of three independent experiments.

**Figure S4 (related to Figure 4).**
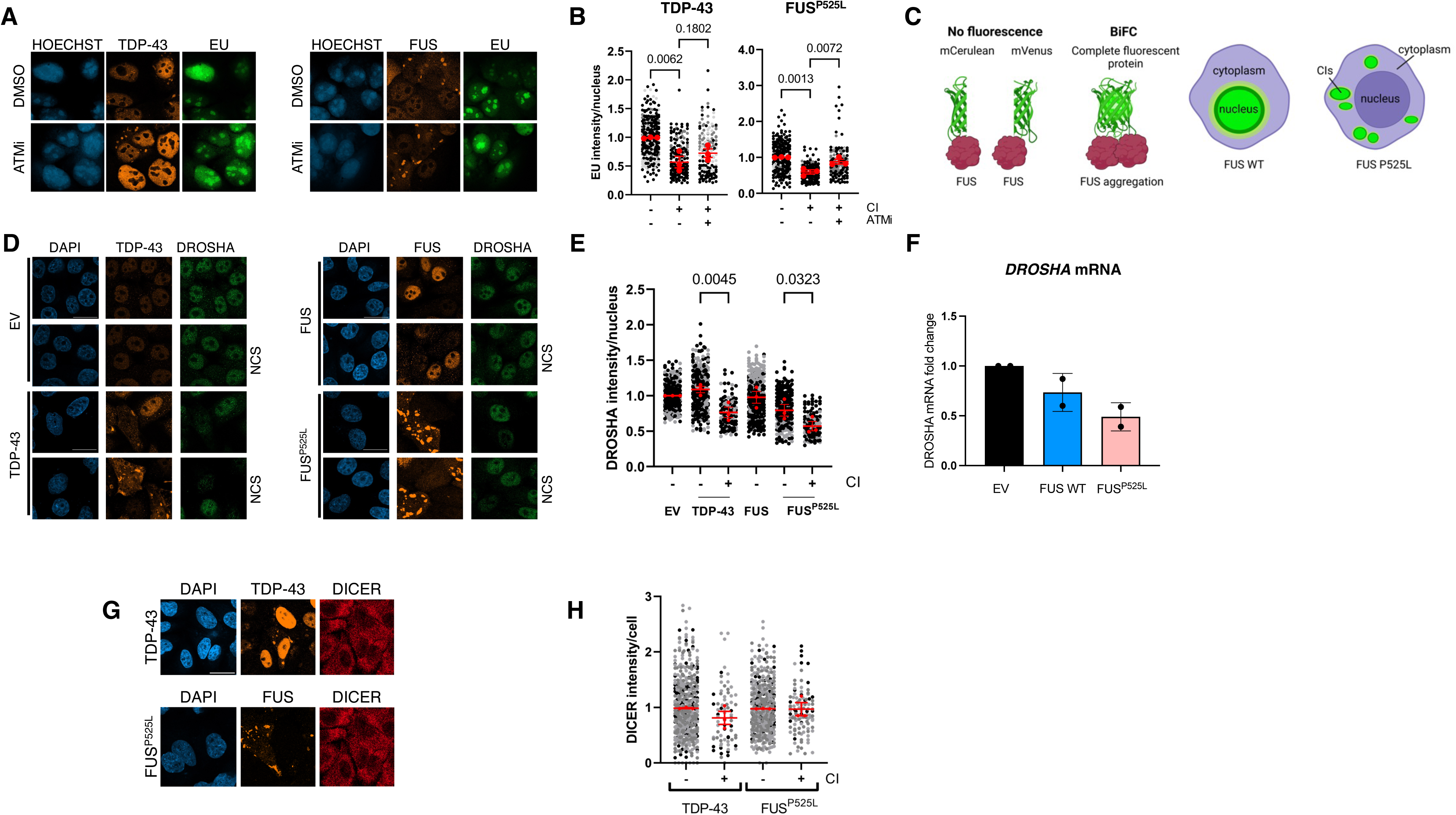
**A)** Representative images of HeLa cells expressing TDP-43 or FUS^P525L^, treated with ATMi or with the vehicle only (DMSO) and pulse-labelled with EU; nuclear DNA was visualized using Hoechst dye. **B)** Quantification of red dots of the super-plots represent the mean values of each biological replicate; red bars indicate the average ± SEM of three independent experiments. **C)** Scheme depicting the Bimolecular fluorescence complementation (BiFC) system used. **D)** Representative images of DROSHA expression in damaged (NCS) or undamaged HeLa cells transfected with plasmids encoding for TDP-43, FUS or FUS^P525L^; cells transfected with an empty vector (EV) were used as a control; nuclei were counter-stained with DAPI; scale bar = 10 µm. **E)** Quantification of DROSHA protein levels in cells with or without cytoplasmic inclusions (± CI) determined in D. The red dots of the super-plot represent the mean values of each biological replicate; red bars indicate the average ± SEM of three independent experiments. **F)** *DROSHA* mRNA levels were monitored by RT-qPCR in HeLa cells selected for the presence of FUS or FUS^P525L^ CIs through FACS. Values are the averages ± SEM of two independent experiments. **G)** Representative images of DICER protein levels in HeLa cells expressing TDP-43 or FUS^P525L^; nuclei were counter-stained with DAPI; scale bar = 20 µm. **H)** Quantification of DICER signals in cells with or without cytoplasmic inclusions (± CI) determined in G. The red dots of the super-plot represent the mean values of each biological replicate; red bars indicate the average ± SEM of three independent experiments.

**Figure S5 (related to Figure 5).**
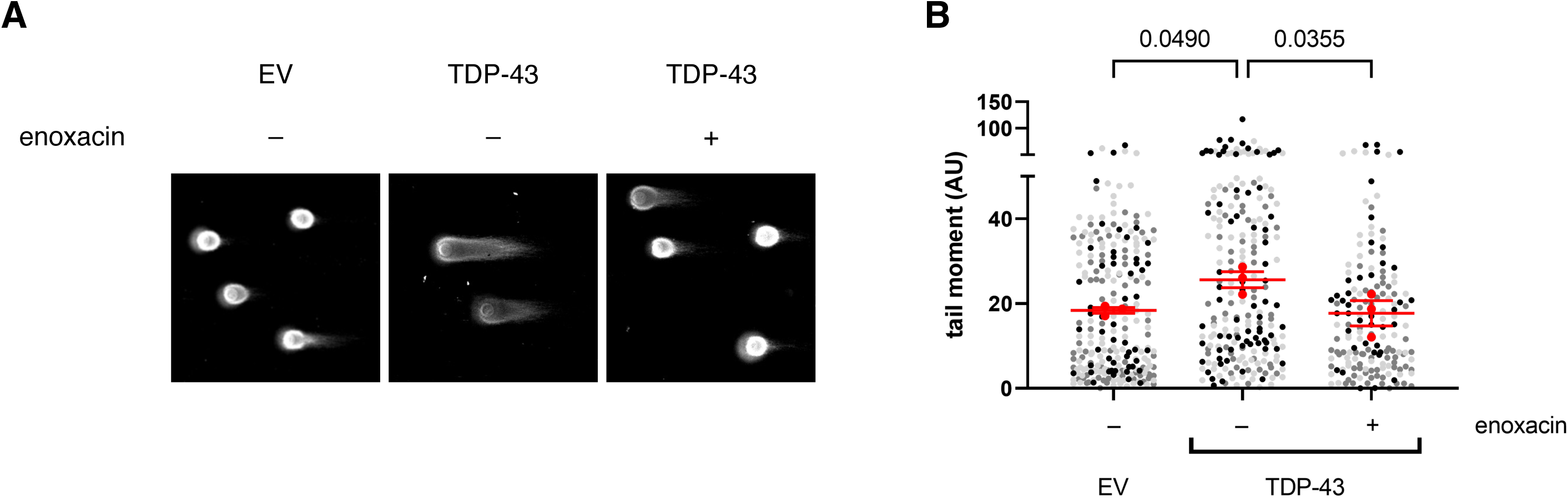
**A)** Representative images of comet assay experiments performed in undamaged HeLa cells transfected with the plasmid expressing TDP-43, or an empty vector (EV) and treated or not with enoxacin. **B)** Tail moment analysis of HeLa cells shown in A. The red dots of the super-plot represent the mean values of each biological replicate; red bars indicate the average ± SEM of three independent experiments.

## Notes

### Competing Interest Statement

The authors have declared no competing interest.

